# Assessing the polymer coil-globule state from the very first spectral modes

**DOI:** 10.1101/2021.07.17.452647

**Authors:** Timothy Földes, Antony Lesage, Maria Barbi

## Abstract

The determination of the coil-globule transition of a polymer is generally based on the reconstruction of scaling laws, implying the need for samples from a rather wide range of different polymer lengths *N*. The spectral point of view developed in this work allows for a very parsimonious description of all the aspects of the finite-size coil-globule transition on the basis of the first two Rouse (cosine) modes only, shedding new light on polymer theory. Capturing the relevant configuration path features, the proposed approach enables to determine the state of a polymer without the need of any information about the polymer length or interaction strength. Importantly, we propose an experimental implementation of our analysis that can be easily performed with modern fluorescent imaging techniques, and would allow differentiation of coil or globule conformations by simply recording the positions of three discernible loci on the polymer.

---

The coil-globule phase transition is a characteristic of the physics of polymers in solution, with direct consequences on macroscopic solution properties as viscoelasticity and transport features. This framework also successfully applies to several biological processes, from protein denaturation to local chromatin density, in which biopolymers compactness abruptly changes under environmental changes. Its widespread applications and deep theoretical relevance justify the relentless effort of both theorists and experimentalists in its understanding and observation. The coil-globule transition is usually studied by means of the scaling properties of the radius of gyration as a function of the polymer length, or number of monomers *N*. However, this approach requires comparing polymers of different lengths, which is not always possible. Even the determination of the number of monomers *N* is not always easy: typically, this number depends on the Kuhn length of the polymer, hence its bending persistence, which can be challenging in the case of real polymers, and a fortiori for complex biological polymers. Moreover, important deviations from the theoretical scaling are induced due to polymer finite-size [1]. An alternative but less direct approach is the study of polymer structure factor and its scaling law as a function of the wave vector in the large *q* limit [2]. We note however that both the radius of gyration and the structure factor are averages over the inter-monomer distances and, therefore, lose information about the precise configuration of the chain in space, which suggests the interest of having another approach that would preserve this information.

From a different perspective, the dynamical behavior of the polymer can be studied. Typically, the dynamics of single monomers can be compared to theoretical predictions such as those of the Rouse or Zimm models [3, 4]. Now, interestingly, the Rouse model uses a spectral mode decomposition. By noticing that this decomposition applies to any given conformation of the chain, we used it to determine the power spectral density as to inquire equilibrium properties of the polymer, rather than its dynamics. Since it contains in principle all the information on the folding state of the polymer, this representation in the reciprocal space can moreover been exploited in the study of the coil-globule transition. The aim of this paper is to investigate this issue and to see how a spectral representation can facilitate the determination of the polymer state (i.e. its macrostate).

By looking in detail at the spectral behavior of polymers close to the coil-globule transition from on-lattice Monte Carlo simulations, we developed a method to discriminate the folding state of a polymer of a given size *N*, without the need to study the scaling law or to have access to the energy parameter. Our method exploits the information about the spatial path of the polymer, and only relies on *low* spectral modes, i.e. long distance features. As we will show, indeed, the very first modes contain the main information concerning the polymer state. This allows us to propose a parsimonious and efficient method, introducing a new *N*-independent order parameter that provides us with a robust measurement to identify the state of the polymer on the coil to globule scale. Most importantly, based on this analysis, we are able to propose an innovative *experimental* approach to determine the equilibrium state of a polymer from data obtained by fluorescence imaging, provided that a minimum of three equally spaced loci can be discernibly labeled, and we provide instructions on how to use these data.

The Rouse model describes an ideal chain, i.e. a chain with no excluded volume nor attractive interaction. In this case, the radius of gyration scales as the variance of a random walk, ⟨*R*^2^⟩ ~ *N*. If hydrodynamic interactions can be neglected, the chain dynamics can be described by applying the overdamped Langevin equation to a polymer of *N* + 1 monomers, who’s positions are denoted by 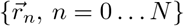, immersed in a fluid of viscosity *γ* at temperature *T* [5]. Subsequent monomers *n* and *n* + 1 are linked by a bonding harmonic potential 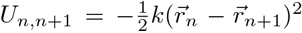, where the spring constant *k* = 3*k_B_T/b*^2^ sets the mean bond length to *b*. The application of the Langevin equation yields a system of *N* + 1 three-dimensional coupled equations of the form

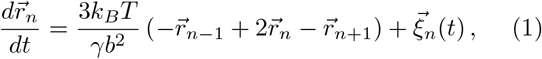

where the random forces *ξ_n_*(*t*), mimicking the effect of thermal fluctuations, satisfy the usual conditions [6]. To solve the model, the system is diagonalized (decoupled) by introducing the new set of variables 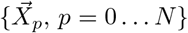

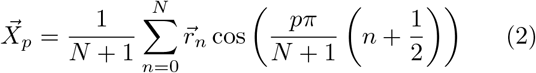

called *Rouse modes* of the polymer and equivalent to the standing modes of an oscillating string with one fixed end. Note that the *p* = 0 mode corresponds to the position of the polymer center of mass. In the following, we will always focus on modes *p* = 1 *… N* only. Usually, one uses the auto-correlation function of 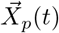 to derive single monomer dynamical scaling laws, overlooking the potentialities of the Rouse decomposition in itself. Indeed, in the language of signal processing, the projection (2) is a Discrete Cosine Transform (DCT), often used for its high performances in image compression. It is closely related to the Discrete Fourier Transform (DFT), in that it corresponds to the DFT of a symmetrized signal of double length [7]. Hence, the mean square mode amplitudes give (up to a multiplicative constant) the *power spectral density* (PSD) of the polymer configuration. For the Cartesian coordinate *x* (and equivalently *y* or *z*) [5]

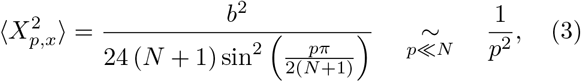

yielding an explicit expression for the PSD of an ideal polymer, and giving an asymptotic power law behavior with exponent – 2.

The connection between Rouse modes and power spectral density seems, surprisingly, to have never been noticed before. However, the asymptotic dependence (3) has an immediate interpretation. Ideal chain configurations are indeed equivalent to Brownian motion trajectories [8], and it can be shown that the corresponding PSD has scaling exponent – 2. It is moreover noteworthy that *exactly* the same expression (3) is obtained in a different setting by Krapf *et al.* for the PSD of a single discrete Brownian trajectory [9].

Now, if the above description holds for an ideal polymer, a real polymer is characterized by interactions between monomers: repulsive excluded volume and (solvent dependent) attractive interactions. The case where only excluded volume is present defines the (non-interacting) *self-avoiding polymer* model, i.e. the prototypical coil polymer. Due to steric hindrance, the polymer occupies more space resulting in swollen, elongated conformations analogous to self-avoiding walks (SAW). The polymer’s radius of gyration then scales as ⟨*R*^2^⟩ ~ *N* ^2*ν*^, where *ν* ≈ 0.588 is the Flory exponent [8]. Including steric interactions adds non-linear terms to the set of equations (1). In this case, the Rouse modes no longer uncouple the equations of motion and, consequently, there is no explicit solution for the PSD [10]. Yet simple arguments provide an insight into the general trend of the spectrum. A usual (yet unproven) assumption about SAWs is their fractal nature, meaning that their statistical properties are scale invariant [8, 11, 12]. Hence, we expect a power law PSD, intuitively related to the presence of long range correlations induced by steric interactions [13]. The Hurst exponent provides an estimation of the long-range memory of a signal. By guessing a long range correlated process with a Hurst exponent *H* such that 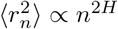, the comparison with the Flory’s behavior would lead to the ansatz *H* = *ν* which appoints the PSD

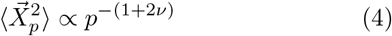

where −(1 + 2*ν*) ≈ −2.2. This is the first prediction to be tested. Interestingly, Panja and Barkema made an equivalent ansatz for the variance of the long wavelengths Rouse modes of the self-avoiding polymer, and corroborate it by extensive on-lattice simulations [10].

Depending on the physico-chemical properties of the polymer and the solvent, an effective attraction *J* between monomers may also exist. The state of the polymer thus depends on the relative strength of inter-monomer forces and thermal energy, *ɛ* = *J/k_B_T*. In the thermodynamic limit *N* → ∞, above a critical value *ɛ_θ_* (*θ*-point), the strong attraction induces a second order phase transition to curled up conformations of uniform density called globules, whose typical scaling is ⟨*R*^2^⟩ ∝ *N* ^2*/*3^ [8]. However, *finite-size* polymers undergo a smooth coil-globule transition at a *N-dependent* critical energy. Due to the fact that there are fewer monomers in *finite-size*, then more energy per contact is required to have globular conformations. Thus, the transition energy 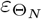 is all the higher that the polymer is small: *ɛ_θ_*(*N*) > *ɛ_θ_*. [2, 14–16].

Coils and globules can therefore in principle be distinguished by their radius of gyration but, as mentioned above, this average quantity does not provide information on the spatial configuration of the polymer, to which the PSD is intrinsically related. The latter can be described by noting with Grosberg and Khokhlov that a collapsed polymer is equivalent to an ideal polymer compressed within a spherical volume of radius *R* [8]. Over small length scales, the polymer behave like a free chain, until it reaches the volume boundary, where it is reflected back and starts an independent random walk. The chain can thus be described as a series of independent subchains. The position of two monomers in different sub-chains is then uncorrelated [8], so that the sequence of positions of monomers sufficiently spaced along the chain is a white noise signal. From the spectral point of view, this implies a constant spectrum for small values of *p*. We can thus predict that the PSD of globules verifies the property

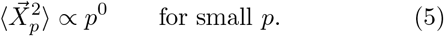

This is the second hypothesis which we now wish to verify.

To test our hypotheses, we simulated single ISAWs cubic lattice [17] and sampled conformations thanks to the Metropolis algorithm with reptation moves [18]. We performed simulations for 22 values of *N* ranging from 100 to 3000, and 100 values of *ɛ* in the range 0.0–0.99. For each (*N, ɛ*), 32000 statistically independent conformations are recorded. Figure 1-a show typical conformations for the case *N* = 431 and three different *ɛ*. The three spatial components are DCT transformed and their square amplitudes are summed to obtain their DCT spectra, as shown in Figure 1-c. For any given (*N, ɛ*), we average over the set of configurations to obtain the corresponding PSD (thick lines in Figure 1-c). Figure 1-d shows the resulting PSDs for different values of *ɛ* and for the given *N*.

**FIG. 1.**
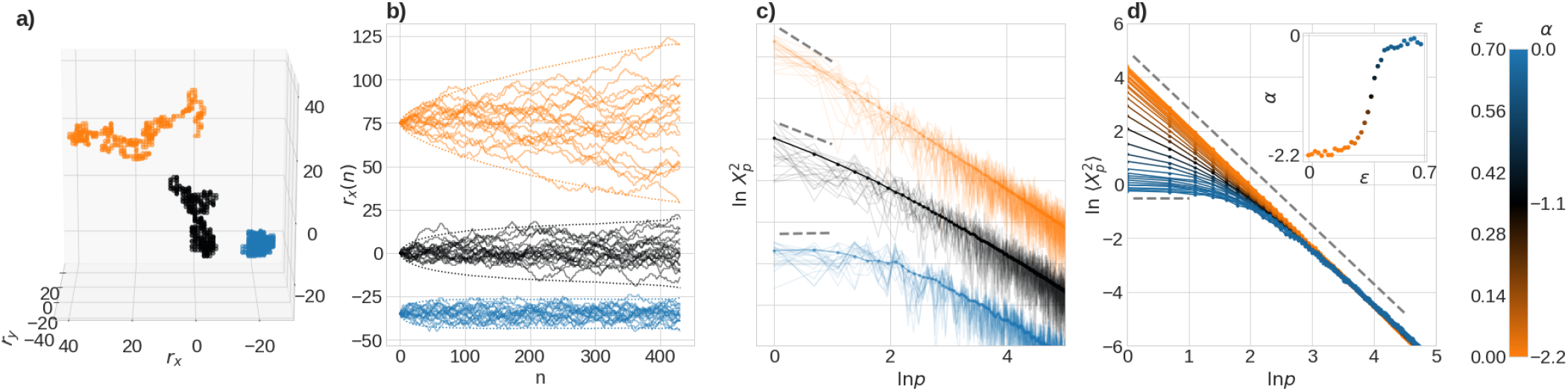
**(a)** On lattice MC simulation snapshots for *N* = 431 and *ɛ* = 0 (orange), *ɛ* = 0.36 (black) and *ɛ* = 0.8 (blue). **(b)** Projections *r_x_*(*n*) along the spatial *x*-axis of a set of 20 configurations, *n* being the monomer index. Colors refer to the same parameters as in (a). Dotted lines correspond to the ±2.5 *σ_x_*(*n*) envelopes. An arbitrary vertical shift is introduced for clarity. (c) Logarithm of the square DCT-transform of the 20 trajectories of (b), ln |*X_p_*^2^|, as functions of ln *p* (thin lines) and their average calculated over the ensemble of 16384 simulated configuration and over the three spatial directions (thick lines and dots). Standard deviations are smaller than the symbol size. Groups of curves are vertically shifted for aim of clarity. The dashed lines correspond to slopes −2.2 ≡ −(1 + 2*ν*), −1.3 and 0 (from top to bottom). **(d)** Averaged PSD obtained by applying the same procedure as in (c) to equivalent sets of simulated polymers with *ɛ* going from 0 to 0.6 (see colorbar). The dashed lines correspond to slopes −(1 + 2*ν*) (top) and 0 (bottom). **Inset:** The slope *α_N_* (*ɛ*) of the log-log representation of the PSD, estimated from the first 2 modes, as a function of the energy parameter *ɛ*.

The results confirm both our predictions. First, for small values of the interaction parameter *ɛ*, the PSDs display the expected scaling of Eq. (4), corresponding to a slope of −(1 + 2*ν*) ≈ −2.2 in the log-log representation. Moreover, the spatial trajectories of Figure 1-b display a typical SAW behavior, with mean square displacement 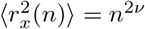.

Second, on the large *ɛ* region, the spatial trajectories clearly show the effect of confinement into the globule volume (Figure 1-b). Correspondingly, the low *p* modes are strongly attenuated, leading to a rather flat spectrum, in agreement with our prediction (5).

Once confirmed our general hypotheses on the spectral profiles for the coil and globule configurations, we address the important issue of how to characterize the coil-globule transition. Inspired by previous observations on the low *p* limit, we defined as a new *order parameter* the log-log slope at *p* = 1 (that is proportional to the logarithm of the ratio of the first two squared modes),

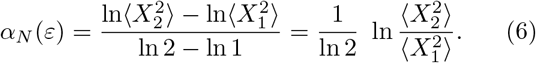

For the case of Figure 1-d, *α_N_* (*ɛ*) clearly performs a *continuous* transition from the typical SAW exponent *α* = −(1 + 2*ν*) to 0, as evidenced in the inset. The width of the transition region depends, however, on *N*. Figure 2 (top panel) shows the *α_N_* (*ɛ*) curves with all the *ɛ* values and for all simulated polymer sizes *N*. These curves have been fitted by a general, 4-parameter sigmoid function, and the fit confirms that all *α_N_* (*ɛ*) interpolates between the two limiting values *α* = −(1 + 2*ν*) (coils) and *α* = 0 (globules). The fact that these limiting values are independent of *N* and *ɛ* makes of *α_N_* (*ɛ*) an excellent order parameter for assessing the polymer state, as it can be applied to each *single* (*ɛ, N*) polymer configuration set and *without the need of any information about these two parameters*. The bottom panel of Figure 2 shows a color plot of *α* as a function of *N* and *ɛ* where colors span from orange, for *α* = −2.2, to blue for *α* = 0. The two regions corresponding to coil and globule states are easily identified, with the typical finite-size crossover region (in black), clearly allowing the definition of a critical transition line *ɛ*(*N*).

**FIG. 2.**
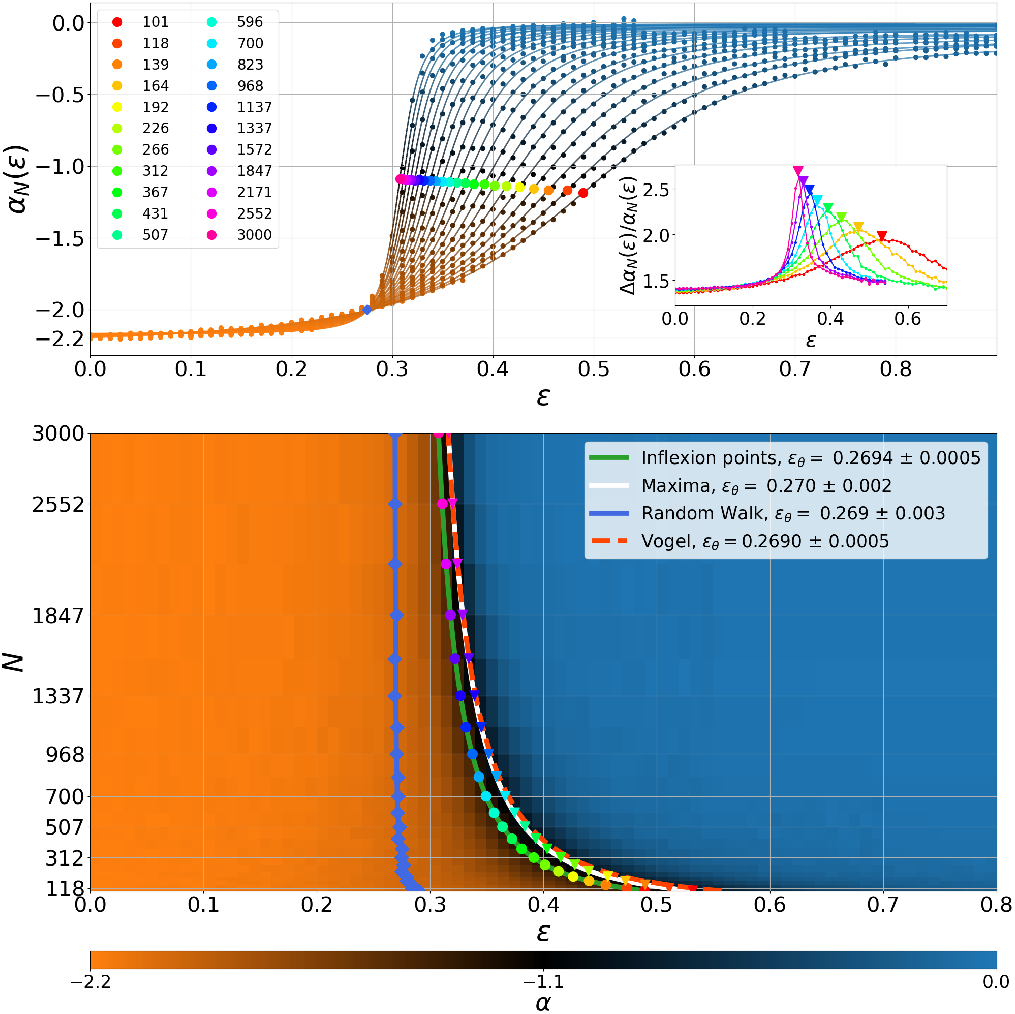
**Top panel:** Log-log plot slope *α_N_* (*ɛ*) estimated on the first 2 modes of PSD (dots) for all polymers sizes *N* from 101 to 3000 (right to left). For each *N*, *α_N_* (*ɛ*) is fitted by a sigmoid 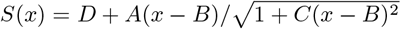 (lines). Colored circles correspond to inflection points, defin-ing *ɛ^I^* (*N*). Blue diamonds indicates the *α_N_* (*ɛ*) = −2 condition, defining 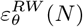. **Inset:***αN* (*ɛ*) relative standard deviation, 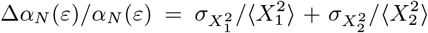, used to define the large fluctuation transition curve. Triangles indicate the position of maxima *ɛ^F^* (*N*), determined by fitting the top of the pics by a Gaussian function. **Bottom panel:** Corresponding phase diagram in the (*ɛ, N*) phase space. The orange to blue colorscale correspond to values of *α_N_* (*ɛ*) from −2.2 to 0 (see colorbar). Inflection points (colored circles) and fluctuation maxima (colored triangles) are shown and fitted by curves of expression *ɛ_θ_*(*N*) = *a*_1_*N* ^−1*/*2^ + *a*_2_(2*N*)^−1^ + *ɛ_θ_* (green and white lines, respectively). The red dashed line is the critical curve of same expression, with the parameters obtained by *Vogel et al.* in Ref. [15]. Fitting parameters are given in Table S1. Both panels: Since the deep globular phase is dominated by low energy, entropically suppressed conformations, a statistical analysis in the high *N*, low *ɛ* region requires sophisticated simulation methods that weren’t undertaken in this study. Samples from the sectors (*N, ɛ*) ∈ [700; 3000] × [0.55; 1.0] and [300; 700] × [0.7; 1.0] didn’t meet statistical relevance and were omitted from the results.

We aimed to compare the scaling of this critical line *ɛ_θ_*(*N*) to previous results on finite-size crossover. Typically, the latter is defined either by the vanishing of the second virial coefficient [14], by the condition of divergent specific heat [15], or by a specific scaling of ⟨*R*^2^⟩ [19]. In Ref. [15], Vogel *et al.* show by on-lattice simulations that the boundary between the two phases is best fitted by a function of the form *f* (*N*) = *a*_1_*N* ^−1*/*2^+*a*_2_(2*N*)^−1^+*ɛ_θ_*. In the framework of our approach based on the first modes, we propose two different definitions for the phase transition critical line. The first one is simply given by the inflection points of the sigmoid functions. The corresponding *ɛ^I^* (*N*) points are very well-fitted by *f* (*N*) (Figure 2, circles, and Table S1). Inspired by the diverging fluctuations typically observed in second order phase transition, we introduce a second, more physical definition of a transition line given by the *ɛ^F^*(*N*) that maximizes the fluctuations of our order parameter *α_N_* (*ɛ*). Indeed, the relative standard deviation Δ*α_N_* (*ɛ*)*/α_N_* (*ɛ*) displays a sharp peak at the transition (meaning that the conformation of the polymer itself largely fluctuates at the crossover) as shown by the inset of Figure 2. The corresponding *ɛ^F^* (*N*) maxima are again best fitted by *f* (*N*) (Figure 2, triangles and Table S1).

Interestingly, both *ɛ^I^* (*N*) and *ɛ^F^* (*N*) curves correspond to values of *α* close to ~ −1.1 (black region) and are therefore far from *α* = −2, associated with RW-like configurations usually identified with the *θ*-point conditions. Our results show, instead, that these RW-like conditions do not appear at the crossover, but are rather located in the coil region of the phase portrait. More precisely, the condition *α_N_* (*ɛ_θ_*) = −2 is met at about *ɛ* = 0.27, close to the point where all the sigmoids roughly cross each other (the blue diamonds in Figure 2) and indeed corresponds to a PSD decreasing as *p*^−2^ over a wide region (Figure S1). The corresponding 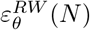 curve is also fitted by a *f* (*N*) function (although with negligible *a*_1_, see Table S1; blue line in Figure 2). As expected, however, the asymptotic *ɛ_θ_* is virtually the same for the three critical curves 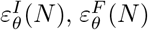 and 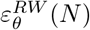 (*N*) and converge, within the error bar, to the well established thermodynamic limit value of *ɛ_θ_* = 0.2690, in agreement with previous numerical estimates [14, 15].

These results point out the richness of the coil-globule transition in finite-size: we were able to distinguish conditions where the chain configurations are RW-like from the thermodynamic (fluctuation based) transition in finite-size. The existence of two distinct critical lines were already pointed out by des Cloizeaux and Jannink [20], but not clearly illustrated by numerical experiments, to the best of our knowledge. As a corollary message of our work, we confirm therefore that the conformation of a polymer of finite-size at the coil-globule transition is not assimilable to a pure random walk. The detailed study of the chain configuration in between these limits, and more generally of their distribution, remains an open question that we will address in an incoming work.

As a last, but most important point, we now come back to the fact that, thanks to the universal values of the order parameter *α*, previous findings allows to assess the state of a single polymer from a sampling of its conformations. Moreover, since this order parameter can be calculated on the basis of the first two spectral modes only, it opens important perspectives on the possibility of *experimentally* assessing the state of a polymer from a very reduced and rather accessible information. In order to get access to the first *M* modes, indeed, it is in principle sufficient to record configurations of *M distinguishable, equally spaced monomers* covering the whole chain, either by following their dynamics or by averaging over a collection of identical polymers to ensure enough statistics. The design of multicolor fluorescent imaging of DNA sites with high spatial and temporal resolution now makes it possible to record the trajectories and relative positions of multiple loci simultaneously [21–23]. If the DCT-transform of a set of *M*-point configurations contain the equivalent of the first modes for the whole chain as we predict, this yields a new experimental approach for the determination of the state of single polymers and biopolymers.

We tested this idea on *decimated* polymer chains, i.e. reduced signals where the positions 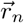 of only *M* equally spaced monomers are retained. We decimated down to *M* = 3, an extreme condition of particular interest from the experimental point of view. Figure 3-a shows examples of PSD obtained for different choices of *M*. Extreme decimation (*M* < 10) alters the first modes. As a consequence, the asymptotic values of the sigmoid *α_N,M_* (*ɛ*) corresponding to “pure” coil (*α_C_*) and globule (*α_G_*) states vary (while, interestingly, the inflection point *α_I_* seems to stay roughly constant), so that it is necessary to provide reference values for these limit slopes as a function of *M*. We give these values in Figure 3-b and in the Table S2. Despite this effect, the asymptotic values for coils and globules remain well apart down to *M* = 3 (Figure 3-b), even for relatively low statistical sampling as shown by the shaded areas, giving a measure of the expected confidence interval on *α* for different sample sizes.

**FIG. 3.**
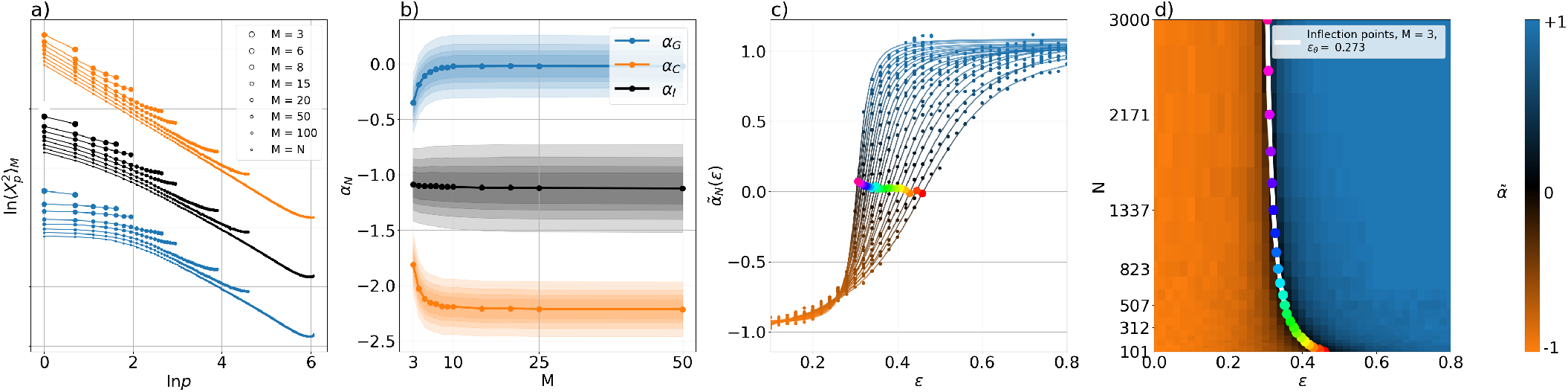
**(a)** PSD obtained by successive decimations for a *N* = 431 chain starting from conformation sets at *ɛ* = 0.0 (orange, coils), *ɛ* = 0.32 (black) and *ɛ* = 0.46 (blue, globules). Decimation goes from *M* = *N* to *M* = 3 (longer to shorter curves). **(b)** The asymptotic and critical values of *α_N,M_* obtained from the sigmoid fit of the decimated spectra, as a function of *M* (dots and lines): *α_C_* (*M*) = *α_N,M_* (0) for coils, orange; *α_I_* (*M*) = *α_N,M_* (*ɛ^I^*) for inflexion points, in black; *α_G_*(*M*) = *α_N,M_* (*ɛ* → ∞) for globules, in blue. These values are averaged over *N*. Shaded regions correspond to 2*σ_α_* confidence intervals for different sample size from 256 (larger intervals) to 2048. They are calculated by propagating the statistical variance of the first two modes through Eq. (6). **(c)** Renormalized sigmoids 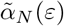 (Equation 7) obtained for *N* from 101 to 3000 with a maximal decimation level *M* = 3. (d) Phase portrait as reconstructed from a *M* = 3 decimated signal. Red dots are obtained from inflexion points as in Figure 2. The blue line is a fit with the same *ɛ_θ_*(*N*) expression, yielding *ϵ_θ_* = 0.268 (see Table S3 for further details). In all graphs we used a dataset of 16384 configurations for each (*N, ɛ, M*) condition.

Once these *M*-dependent limit values obtained, the order parameter *α_N,M_* (*ɛ*) can be normalized as

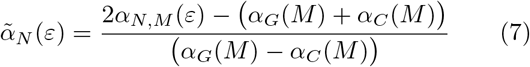

so to span from −1 to 1. In this way, equivalent sigmoids 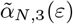 are independent of *M* (Figure S2). For sufficiently large samples, even an extreme decimation with only 3 equally spaced points allows a very accurate reconstruction of the (*N, ɛ*) phase diagram, as shown in Figure 3-d. The critical line *ɛ^I^* (*N*) matches that of the complete chain (white line in Figure 3-d and Table S3).

We finally checked the robustness of the proposed approach against variations in the size of the statistical sample. The obtained phase portraits are shown in Figure S3. We have been able to reconstruct the entire phase diagram from the configurations of only three points, by exploring the (*N, ɛ*) parameter space. Obviously, trying to determine the state of a polymer under given conditions, without being able to vary any parameter, is further complicated by the fact that finite-size effects generate a relatively large crossover region in which the state of the polymer smoothly change from coil to globule. However, the great robustness of the method introduced in this work and the universal character of the 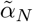 parameter allow us to conclude that the proposed approach potentially outperforms any other method aiming at determining in which phase a polymer in given conditions falls. An important additional advantage comes from the fact that our approach does not require knowledge of the length of the polymer in terms of Kuhn lengths, nor of the magnitude of the attraction between monomers, a parameter otherwise difficult to characterize.

The three-point measurement is very easy to perform experimentally, especially since, in this case, it is sufficient to discern the central point from the two extremities, which can be done for example by two-color fluorescence. Finally, preliminary results indicate that the method is also extremely robust to variations in the position of the three selected monomers. To go even further, we will implement an upgraded method that allows to explicitly take into account their position, in order to obtain results as good as for perfectly equidistant monomers. This improvement will be presented in a future work.

The authors would like to acknowledge networking support by the EUTOPIA COST Action CA17139. This work has been partially supported by the ANR project ANR-19-CE45-0016. Vincent Dahirel and Jean-Marc Victor are warmly acknowledged for their constant support and insightful advice.

## Supplementary informations

### Coefficients *a*_1_, *a*_2_ and *ɛ_θ_* for the three curves

**TABLE S1.** In Figure 2 we have fitted inflection points (colored circles) and *α_N_* (*ɛ*) fluctuation maxima (colored triangles) by the expression *ɛ_θ_*(*N*) = *a*_1_*N* ^−1*/*2^ + *a*_2_(2*N*)^−1^ + *ɛ_θ_* from Ref. [15] (green and white lines, respectively). Here we list the corresponding fitting parameters *a*_1_, *a*_2_ and *ɛ_θ_*, together to those obtained by Vogel *et al.* in Ref. [15] by fitting a different order parameter.

| Data | $a_1$ | $a_2$ | $\varepsilon_\theta$ |
| --- | --- | --- | --- |
| Inflection points $\varepsilon^I(N)$ | $2.1 \pm 0.04$ | $3.3 \pm 0.9$ | $0.2694 \pm 0.0005$ |
| Fluctuation maxima $\varepsilon^F(N)$ | $2.5 \pm 0.09$ | $2.7 \pm 1.8$ | $0.270 \pm 0.002$ |
| Random Walk $\varepsilon^{RW}(N)$ | $0.0 \pm 0.1$ | $4.6 \pm 1.8$ | $0.269 \pm 0.003$ |
| Vogel *et al.* $dC_v/dt$ [15] | 2.5 | 8.0 | $0.2690 \pm 0.0005$ |

### Reference values *α_C_*, *α_I_* and *α_G_* obtained for different *M*

**TABLE S2.** Numerical values of the loglog-slopes *αN,M* (*ɛ*) obtained from the spectra in Figure 3-a and displayed in Figure 3-b (points), as a function of *M*. The parameter *α_C_*, for coils, is obtained at the limit *ɛ* = 0; *α_I_* is measured at the fitting sigmoid inflexion point; *α_G_*, for globules, is calculted as *ɛ* → ∞ limit of the fitting sigmoid.

| $M$ | $\alpha_C^a$ | $\alpha_I^a$ | $\alpha_G^a$ |
| --- | --- | --- | --- |
| 3 | -1.81 | -1.09 | -0.36 |
| 4 | -2.03 | -1.10 | -0.19 |
| 5 | -2.11 | -1.10 | -0.11 |
| 6 | -2.15 | -1.10 | -0.07 |
| 7 | -2.16 | -1.10 | -0.05 |
| 8 | -2.19 | -1.11 | -0.03 |
| 10 | -2.19 | -1.11 | -0.03 |
| 20 | -2.21 | -1.12 | -0.02 |

<sup>a</sup> Significant digits.

### Critical *ɛ^I^*(*N*) inflection point fit parameters upon decimation

**TABLE S3.** As shown in Figure 3, decimated configurations retain the main information about the polymer state and allows us to reconstruct the phase diagram in the (*ɛ, N*) phase space. Moreover, inflection points (colored circles in Figure 3) can be used to define a critical line also in decimated conditions, and fitted by the function *ɛ_θ_*(*N*) = *a*_1_*N* ^−1*/*2^ + *a*_2_(2*N*)^−1^ + *ɛ_θ_*. Here we give corresponding values of the three fitting parameters for *M* = 3, 5, 8, 10 and, for comparison, for the non decimated case *M* = *N*. Note that the asymptotic value *ɛ_θ_* is essentially independent on decimation, ensuring a correct definition of the phase transition in the thermodynamic limit. The *a*_1_ and *a*_2_ parameters, that gives the *N* dependence of the transition in finite-size, are instead sligthly modified by the decimation procedure, but converge toward the original values as soon as *M* ≳ 10.

| $M$ | $a_1$ | $a_2$ | $\varepsilon_\theta$ |
| --- | --- | --- | --- |
| 3 | $1.7 \pm 0.1$ | $2.7 \pm 1.8$ | $0.273 \pm 0.002$ |
| 5 | $1.8 \pm 0.05$ | $5.0 \pm 0.9$ | $2.270 \pm 0.001$ |
| 8 | $1.9 \pm 0.04$ | $4.4 \pm 0.7$ | $2.270 \pm 0.001$ |
| 10 | $2.0 \pm 0.05$ | $4.2 \pm 0.8$ | $0.270 \pm 0.001$ |
| $M = N$ | $2.1 \pm 0.04$ | $3.3 \pm 0.9$ | $0.2694 \pm 0.0005$ |

### Typical PSD at the theta conditions 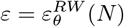

**FIG. S1.**
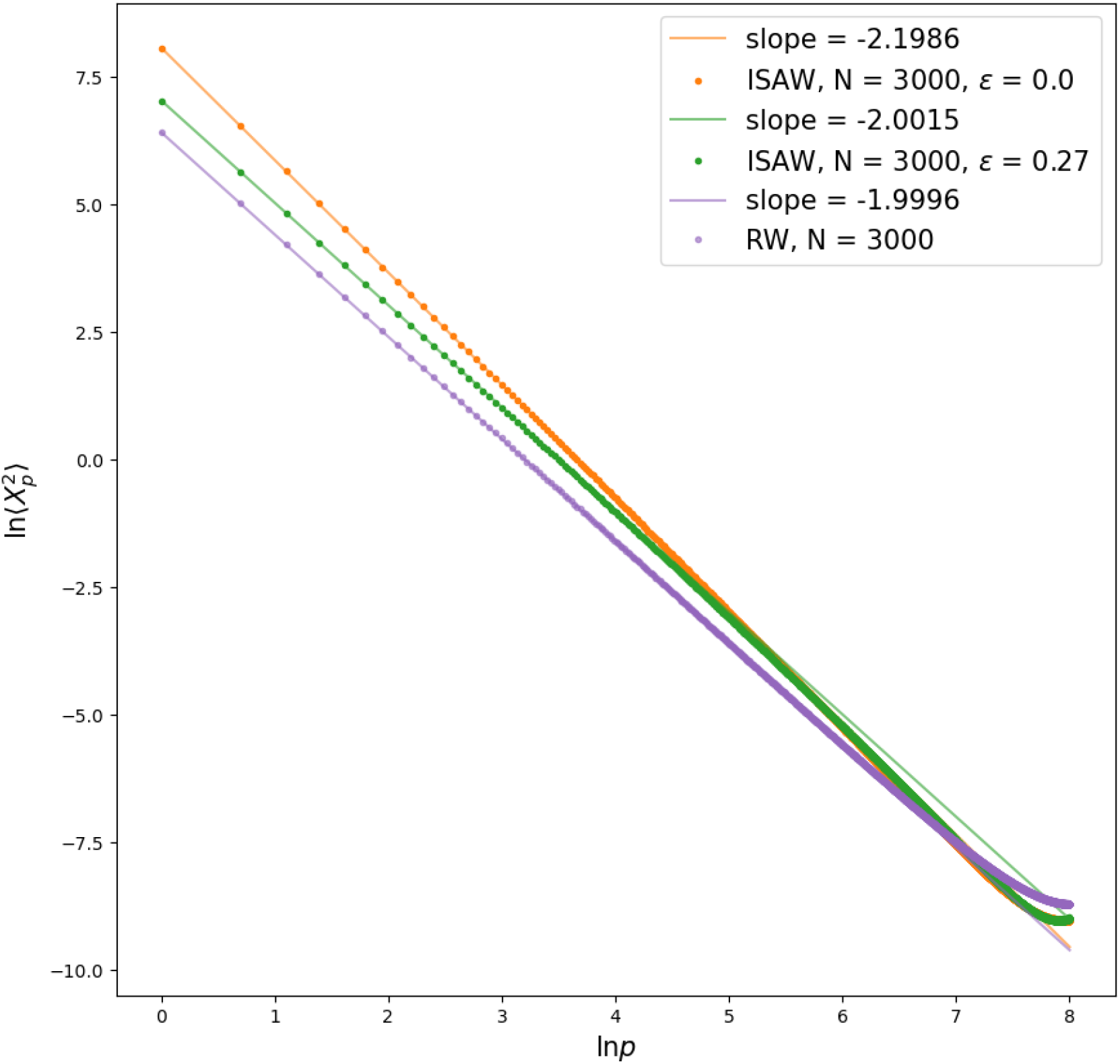
The plot shows a typical PSD obtained for a value of *ɛ* matching the *α* = −2 conditions, i.e for 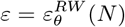 (green points). For comparison, a pure coil spectrum is shown (orange points) with its typical −2.2 slope (orange line) and a pure on-lattice RW (purple points) of slope −2 (purple line) are shown. The first modes slope *α* for 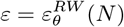 is as expected equal to −2 (green line); Moreover, the plot shows how the same scaling law holds over a wide range of modes. This ensures that the polymer configurations can be described as random walks up to short scales (where a SAW behavior is recovered).

### Effect of different levels of decimation on the phase transition

**FIG. S2.**
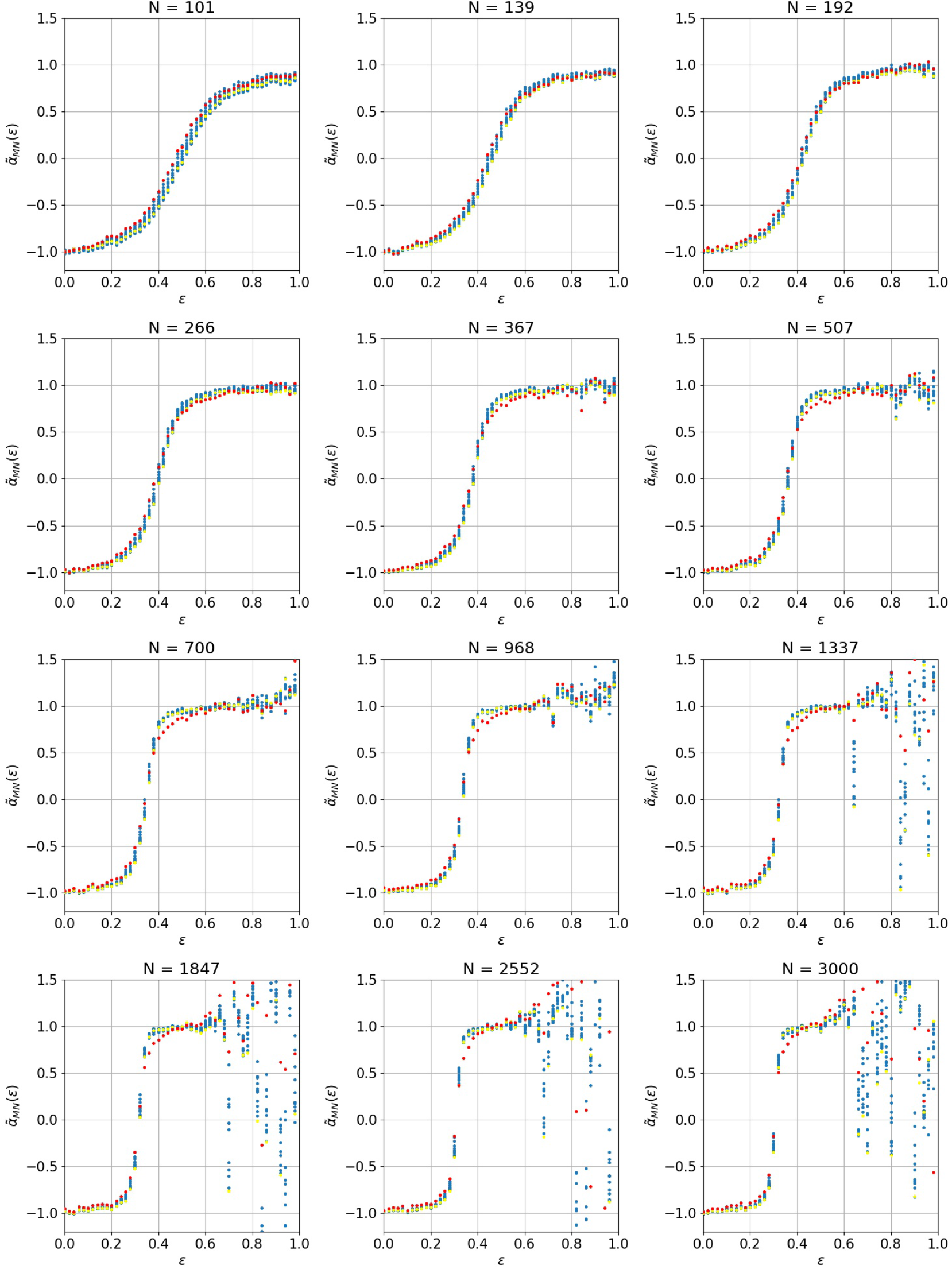
The different panels show renormalized loglog-slopes 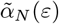 (Eq. (7)) for different value of *N*. For each *N*, we show results obtained by different levels of decimation *M* (blue points), starting from 3 (red points) to *N* (yellow points). Comparison of these different cases shows that, for given *N*, there is essentially no difference in the behavior of the order parameter *α_N_* (*ɛ*), once rescaled with the maximum and minimum values, *α_G_*(*M*) and *α_C_* (*M*), respectively. Data sample includes here 16384 configurations per point. We recall that, as in Figure 2, samples from the sectors (*N, ɛ*) ∈ [700; 3000] × [0.55; 1.0] and [300; 700] × [0.7; 1.0] didn’t meet statistical relevance. Here we have kept the raw results from these sectors, which also gives a measure of the variability of the corresponding values.

### Phase portraits after decimation, with different sample size

**FIG. S3.**
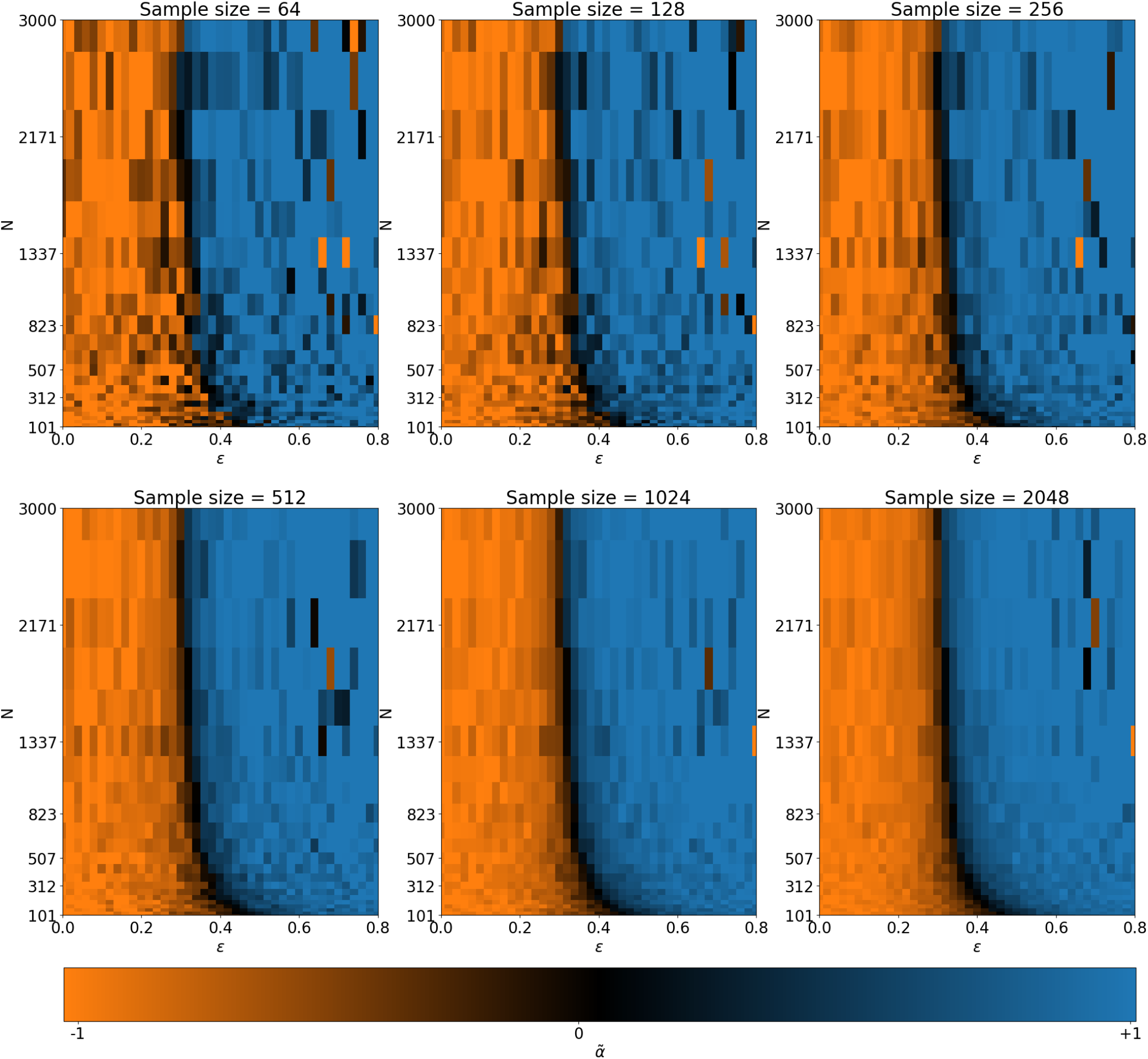
Example of phase portraits obtained by extreme decimation *M* = 3 starting from a same set of conformations, for different *sample* sizes, from 64 to 2048. We recall that, as in Figure 2, samples from the sectors (*N, ɛ*) ∈ [700; 3000] × [0.55; 1.0] and [300; 700] × [0.7; 1.0] didn’t meet statistical relevance. Here we have kept the raw results from these sectors, which also gives a measure of the variability of the corresponding values. Overall, we observe that if a sample of only 64 configurations is not sufficient to assign with certainty a system to the coil or globule states, it becomes rather good starting from 512.

